# Expectation modulates learning emotional words: Evidence from a hierarchical Bayesian model

**DOI:** 10.1101/2024.07.25.605051

**Authors:** Weiwei Zhang, Yingyu Li, Chuan Zhou, Baike Li, John W. Schwieter, Huanhuan Liu, Meng Liu

## Abstract

In language acquisition, individuals learn the emotional value of words through external feedback. Previous studies have used emotional words as experimental materials to explore the cognitive mechanism underlying emotional language processing, but have failed to recognize that languages are acquired in a changing environment. To this end, this study aims to combine reinforcement learning with emotional word learning, using a probabilistic reversal learning task to explore how individuals acquire the valence of emotional words in a dynamically changing environment. Our computational modeling on both behavioral and event-related potential (ERP) data revealed that individuals’ expectations can modulate the learning speed and temporal processing of emotional words, demonstrating a clear negative bias. Specifically, as the expected value increases, individuals respond faster and exhibit higher amplitudes for negative emotional words. These findings shed light on the neural mechanisms of emotional word learning in a volatile environment, highlighting the crucial role of expectations in this process and the preference for processing negative information.

## 1. Introduction

For individuals to better adapt to survive and develop, the ability to quickly and efficiently process information containing emotions is crucial (Damasio, 2001). Language serves as an important element of human survival and development, and plays an important role in conveying diverse emotions. Individuals acquire the emotional value of words through positive or negative feedback from the outside world in a dynamically changing environment (Albrecht et al., 2023; Frishkoff et al., 2016; Goldstein & Schwade, 2008; Pashler et al., 2005; Schindler et al., 2019; Scott et al., 2009; Snefjella et al., 2020). Currently, a growing number of studies are utilizing emotional words as experimental materials to investigate the mechanisms underlying emotional language processing (Wu & Zhang, 2019; Zhang, Teo, & Wu, 2019; Zhang et al., 2017, 2019, 2020). However, in real life, the learning of emotional words takes place in a dynamically changing environment. In the present study, we utilize the principles of reinforcement learning to simulate a dynamically changing environment in order to investigate how humans acquire the valence of emotional words.

Over the past half century, reinforcement learning (RL) has become one of the most widely used computational models in cognitive psychology and neuroscience (Hackel & Amodio, 2018). It is often used to solve problems of uncertainty in decision-making and various learning-related issues (Diaconescu et al., 2014; Den et al., 2013; Lefebvre et al., 2017; Lee et al., 2012; Mukherjee et al., 2020; Palminteri & Lebreton, 2022; Sharp et al., 2022). Regarding the learning and decision-making processes in dynamic environments, researchers have proposed a more flexible, efficient reinforcement learning model known as the Hierarchical Gaussian Filter (HGF) model (Mathys et al., 2011; Mathys et al., 2014). The HGF model integrates reinforcement learning with Bayesian learning, allowing it to mathematically quantify the internal learning process of an individual and to infer the trajectory of belief updates, effectively revealing individuals’ learning process in dynamic and uncertain environments. The most commonly used experimental paradigm in reinforcement learning is the probabilistic reversal learning task (Browning et al., 2015; De Berker et al., 2016; Liu et al., 2022; Liu et al., 2023; Metha et al., 2020; Mukherjee et al., 2020). In this paradigm, the relationship between responses and outcomes can be reversed throughout the entire experimental process (Diaconescu et al., 2014; Hein et al., 2021; Hein & Ruiz, 2022; Liu et al., 2022), creating a fluctuating environment by manipulating different probabilities. Within this hierarchical learning framework, expectation is an essential concept. The model posits that in a constantly changing environment, an individual will use external feedback to continuously update their expectations about the probability of receiving a reward for given choices, as well as their expectations about the rate of change of such a probability. This enables an individual to correctly learn the appropriate rules and flexibly adapt to the environment. In this process, expectations play a central role in shaping and influencing behaviour (Hein et al., 2023; Hein & Ruiz, 2022). With respect to language learning, external feedback, such as encouragement and criticism from other individuals, affects learners’ expectations of the outcome of the language expressed. For example, when individuals speak words that conform to social norms or express emotions appropriately in different social situations, they may have expectations about how their expressions will be perceived. Regardless of whether the actual outcome meets their expectations, individuals continuously accumulate experience and form prior expectations from their experience. These expectations subsequently guide their successive behaviours. Previous researchers have combined reinforcement learning with studies in other fields (Li et al., 2023; Liu et al., 2023; Hackel et al., 2020; Held et al., 2024; Lockwood & Klein-Flügge, 2021; Wang et al., 2023; Zhang et al., 2023). For instance, Wang et al. applied reinforcement learning to emotion processing and elaborated on the cognitive neuroscientific mechanisms of how fear interferes with adaptation to environmental volatility in dynamic environments at the computational level. However, in the field of linguistic research, fewer studies have investigated the impact of extra-linguistic factors on language processing. In the present study, we combine reinforcement learning with emotional word learning using the HGF model to explore the acquisition process of emotional words in a simulated authentic and natural scenario.

As individuals continuously interact in their external environments, they gradually associate specific words with emotional experiences through feedback, thereby acquiring the valence of different emotional words (Albrecht et al., 2023; Frishkoff et al., 2016; Pashler et al., 2005). The emotional values of these words, also known as valence, expresses the positive or negative attributes associated with them (Kauschke et al., 2019; Russell, 1980). However, research has found that valence bias exists in the learning of emotional words (Altarriba & Basnight-Brown, 2011; Degner et al., 2012; Eilola et al., 2007; Estes & Verges, 2008; Nasrallah et al., 2009; Sutton et al., 2007; Wu et al., 2022; Zhang, Teo, & Wu, 2019). On the one hand, some studies have found that individuals have a positive bias in processing emotional words, with some results indicating that individuals recognize positive words faster than negative words (Eilola & Havelka, 2011; Hinojosa et al., 2015; Kuperman et al., 2014; Ponari et al., 2015; Schindler & Kissler, 2016; Sylvester et al., 2016; Yao et al., 2016) and have an advantage in accuracy when processing positive words (Bayer & Schacha, 2014). The density hypothesis proposed by Unkelbach et al. (2008) provides a detailed explanation for the asymmetry in the processing speed of positive and negative stimuli. The hypothesis suggests that positive information is stored with a higher density compared to negative information, resulting in positive stimuli being more quickly processed than negative stimuli. Zhang, Wu, Yuan and Meng (2019) expanded on the hypothesis, arguing that positive emotion-label words are clustered in density, while negative emotion-label words are stored discretely, leading to faster processing speeds for positive emotion-label words. On the other hand, some studies have also found that individuals have a negative bias when learning emotional words (Estes & Verges, 2008; Nasrallah et al., 2009). For instance, Zhang et al. (2017) revealed that negative emotion-label words elicited stronger late positive component (LPC) effects than positive emotion-label words and showed a right-sidedness advantage. Furthermore, Wu et al. (2022) demonstrated that negative emotion-label words can facilitate individuals’ perception of emotion for subsequent stimuli. The positivity-negativity asymmetry hypothesis proposed by Robinson-Riegler and Winton (1996) suggests that negative stimuli have a higher emotional intensity than positive stimuli, leading individuals to be more inclined to process negative stimuli (see also Smith et al., 2003; Taylor, 1991). The automatic vigilance hypothesis provides another explanation for the negative bias (Pratto & John, 1991), such that automatic evaluation of stimuli is a mechanism that directs attention directly to events related to life threats. Therefore, the human brain tends to filter and analyze negative information from the environment to avoid adverse outcomes. However, there are discrepancies between positive or negative biases in emotional word learning, making it worthy of further exploration.

In the present study, we employ a reinforcement learning task to explore how individuals update their expectations and acquire the emotional valence of words through feedback in a dynamically changing environment. We asked participants to identify formation rules of words under two different experimental conditions: positive and negative emotion-label words. In doing so, participants were required to flexibly adapt to changes in the environment and to select words that had the highest probability of obtaining a reward (see Figure1A). For instance, under the positive emotion-label word condition, two types of words (composed of “cu” or “zo” suffixes) were presented to participants. In the stable environment phase, words ending in “cu” consistently had an 80% reward probability, while words ending in “zo” had a 20% reward probability. However, in the volatile environment phase, the reward probabilities for these two sets of words undergo rapid fluctuations, alternating between 80% and 20% for each set five times throughout the phase. Given previous research indicating the existence of positive or negative biases in the learning of emotional words, we use a probabilistic reversal learning task and hypothesize that individuals may also exhibit a preference for either positive or negative emotional words when learning the valence of words in a dynamically changing scenario.

## 2. Methods

### 2.1 Participants

Twenty-eight individuals participated in this study. Six participants were not included in the analysis: two participants did not follow the instructions and another four had excessive EEG artifacts. Therefore, the final sample comprised 22 participants (13 females, 9 males, age range: 18-26 years, age: *M* = 22.05 years, *SD* = 1.94). All participants reported no neurological or psychiatric disorders and had normal or corrected-to-normal vision. The experiment was approved by the Research Center of Brain and Cognitive Neuroscience in Liaoning Normal University, and all volunteers provided their written informed consent prior to participating in the study. All participants were native Chinese speakers with an intermediate level of English proficiency.

### 2.2 Materials

Consistent with a study by Gong et al. (2016), we created 40 pseudowords (20 words for each experimental condition, see supplementary Table S1). All pseudowords were composed of four lowercase letters. For positive emotional words, the pseudowords were divided into two types: those whose suffix was “cu,” and those whose suffix was “zo”. For negative emotional words, the pseudowords had “da” and “ye” suffixes. The environment in the experiment included stable and fluctuating phases. In the stable phase, there was an 80% probability of receiving a reward for words belonging to pseudowords with “cu” and “das” suffixes. However, in the fluctuating phase, the probability of receiving a reward alternated between 20% and 80% for pseudowords with “zo” and “ye” suffixes compared to those with suffixes “cu” and “da”, and this switch occurred five times.

From the Test of Emotion Comprehension (Pons & Harris, 2000; Pons et al., 2004), 66 simplified facial expression line drawings depicting different facial expressions were selected. The complexity, size, and clarity of the pictures, the thickness of the lines, as well as the gender and age of the depicted individuals, were all controlled. Thirty-seven adults who did not participate in the formal experiment were asked to rate the valence and arousal of these images on a nine-point Likert scale. Based on their ratings, 39 line-drawing pictures were selected, with 13 each representing positive, neutral, and negative emotions. A one-way ANOVA revealed significant differences in valence among the three groups of pictures, *F*(2,36)=758.858, *p* < .001, however, no significant difference was found in arousal levels, *F*(2,36) =.118, *p* =.737 (see Table 1). For the formal experiment, one representative picture from the positive and negative categories was selected for further study.

**Table 1.**
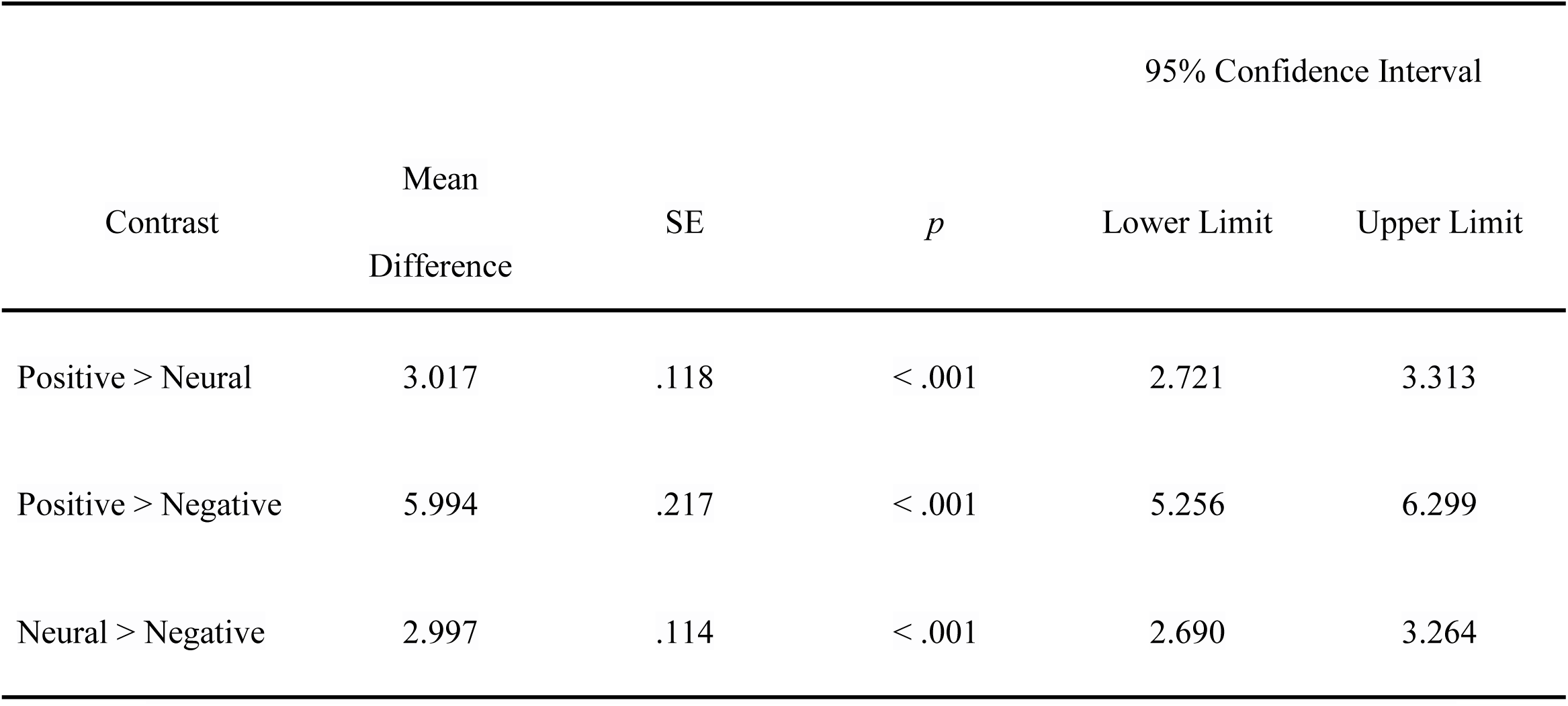
Pairwise comparisons with Bonferroni corrections for the valences of emotion task.

### 2.3 Experiment Procedure

We employed a within-subject experimental design using emotional word type (positive and negative words), which was implemented in E-prime 2.0. All participants completed the task consisting of two blocks of 100 trials each: block 1 (positive emotional words), block 2 (negative emotional words). Each block lasted approximately 12 minutes and participants were given a brief break between the two blocks.

Prior to the formal experiment, we explicitly informed participants that the reward structure would be changed throughout the entire task (De Berker et al., 2016). We asked them to first practice the task with stimuli which differed from the formal experiment to familiarize themselves with the procedure. In each block, participants were asked to complete a probabilistic reversal learning task. The goal was to find the association between formation rules (positive or negative emotional words) and correct feedback. Thus, they had to learn the probability of the rewards assigned to each association. Each block comprised two environments with different association contingency mappings: in the stable environment (50 trials), the probability of reward for correct association was .8; in the volatile environment, this part was divided into 5 subsections (50 trials,10 trials for each subsection), and the probability of reward for correct associations switched between .2 and .8 on five occasions (see Figure 1A).

**Figure 1.**
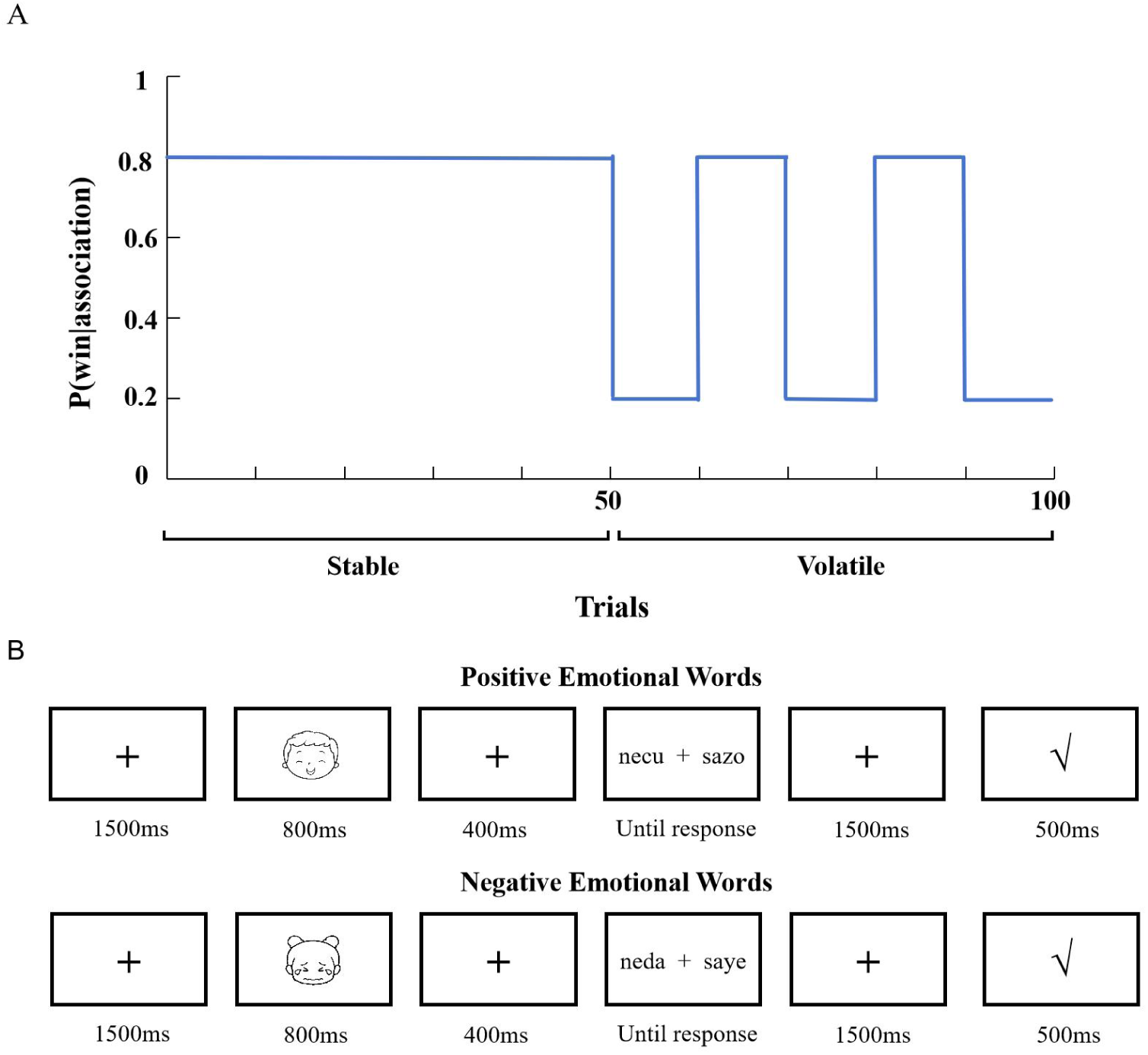
Experimental design exploring the impact of a fluctuating reinforcement learning environment on the learning of positive and negative emotional words. A) The probability settings for correct association. B) Examples of trial procedures for the conditions of positive and negative emotional words.

Figure 1B shows the procedure of the formal experiment: each trial began with a fixation cross for 1500 ms followed by the presentation of an emotional picture for 800 ms. After a 400 ms fixation cross, participants were asked to identify which word would receive correct feedback and responded (press ‘F’ or ‘J’). Only after subjects made their choice, the outcome, either with correct or wrong feedback, was shown on the screen for 500 ms after a fixation cross appeared for 1500 ms. The participants were randomly assigned to either complete the condition for positive emotional words first, or the condition for negative emotional words first. Similarly, half of participants performed the stable environment first and the other half of participants performed the volatile environment first.

### 2.4 HGF Model

For the current binary probabilistic reversal learning task, we used the Hierarchical Gaussian Filter (HGF) Model established by Mathys et al. (2011, 2014) to estimate each participant’s belief trajectories. We used a 3-level HGF model to fit the current behavioral results. Figure 2 illustrates the perception model and response model.

**Figure 2.**
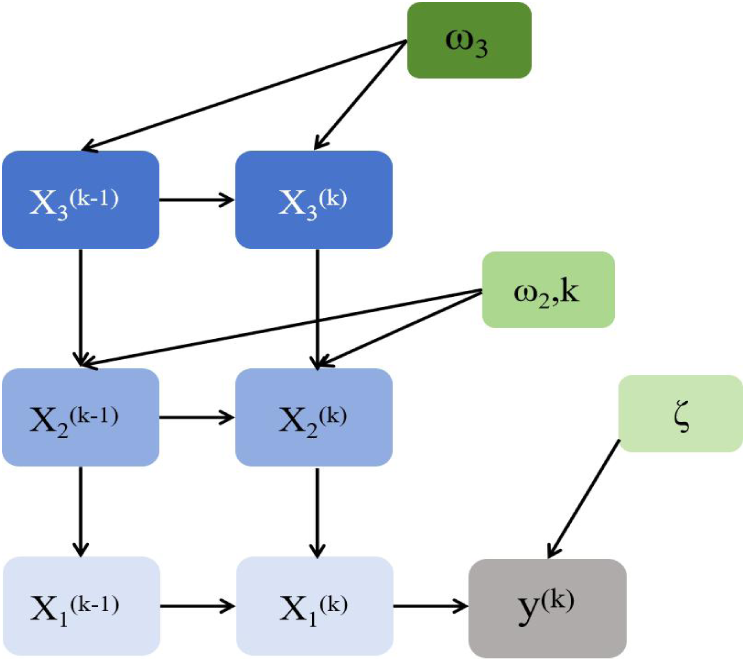
Schematic representation of the 3-level HGF model. *Note. ω*2, *ω*3 and *ζ* were fitted, and *k* was fixed to 1.

#### 2.4.1 Perceptual Model

The perceptual model is divided into three layers with corresponding hidden states (x_1_, x_2_, x_3_). At the first level, x_1_ is a discrete variable, which represents the binary stimulus category: whether a word with a certain rule type (e.g., suffix cu/da) is rewarded (x_1_(k) = 1) or not (x_1_(k) = 0) in trial *k*, while the other type of word (e.g., suffix zo/ye) is not rewarded or rewarded accordingly. The states in levels two and three are continuous variables: x_2_ corresponds to participant’s tendency towards words composed of cu/da suffixes and x_3_ represents the log-volatility of tendency on the second level. Because the probability distributions of the second and third layers are evolving as Gaussian random walks, their value at trial *k* will be normally distributed around its previous value at trial *k-1.* At the two levels, the mean and variance of posterior distribution of beliefs about x_i_ (i = 2, 3) are *μ_i_* (an individual’s expectation) and *σ_i_* (uncertainty).

There are also some parameters that reflect the changes in the states of each level and the interaction between them: *k* represents the coupling strength between 2-level and 3-level, we fixed it to 1 (Hein, de Fockert, & Ruiz, 2021); ω_2_ corresponds to the constant part of the step size of 2-level; ω_3_ is a constant part, which determined the step at the third layer.

For each trial and level *i* (*i* = 2, 3), the beliefs of the posterior means were updated per the following equation: 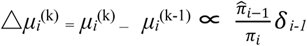. In the equation: △*μ_i_* ^(k)^ represents the posterior mean update, which denotes the difference between the posterior expectation in the current trial *k* and previous trial *k-1*; *δ_i-1_* corresponds to prediction error, which was from a lower layer and was used to update belief in the current level; and 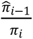 was a precision-weights ratio, in which 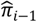 and *π_i_* were the prediction precision of the level below and the precision of the current belief, respectively. The precision can be defined as the inverse variance of the posterior expectation: *_πi_*^(*k*)^ *_=_* _1/*σi*_^(*k*)^. In the current study, our analyses focused on participants’ expectations in the 2-level, corresponding to the parameter *μ_2_*.

#### 2.4.2 Response Model

A response model described how an individual predicted and evaluated the probabilistic consequences of behaviour based on a generated perceptual model, and finally made the most advantageous decision. For binary inputs, we converted predicted reward probabilities m(k) for stimuli (e.g., pseudowords with “cu” suffix) using a unit-square sigmoid model to probabilities of subjects acting on those stimuli (Hein et al., 2021; Hein & Ruiz, 2022; Iglesias et al., 2013; Mathys et al., 2014): 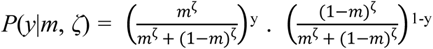. Here, higher values of the temperature parameter *ζ* indicated that individuals were more confident about their choice, i.e., they were more likely to choose stimuli that were consistent with their current beliefs.

### 2.5 Model Space

We compared three computational models of learning and used random effects Bayesian model selection (BMS) to determine the best model (Stephan et al., 2009). The first was a three-level HGF (Mathys et al., 2011, 2014), which incorporates the strengths of reinforcement and Bayesian learning, providing important information about hierarchical learning (Liu et al., 2022). Two additional control models were both non-hierarchical models: the second was Rescorla Wagner (RW), which is one of the most prevalent and simplest models of reinforcement learning, and which assumes that prediction errors drive belief updating, but its learning rate is fixed (Rescorla & Wagner, 1972); and the third was a SuttonK1 model (SK1), which permits learning rate to be adaptive and is affected by prediction errors in the most recent trials (Sutton, 1992). According to three indicators (exceedance probability, protected exceedance probability and model frequencies) provided by BMS (Soch et al., 2016), we chose the winning model for further analyses.

### 2.6 Behavioral Data Analysis

We performed a generalized linear mixed-effects model analysis and a generalized logistic mixed-effects model analysis for RTs and accuracy, respectively (R 4.0.1, lme4 package) (Douglas Bates et al., 2015). Experimental conditions (positive and negative emotional words) and the parameter (*μ_2_*) were added as fixed effects, while participants were added as a random effect. To reduce the influence of non-normally distributed data, RTs of all trials were log-transformed. Consistent with Visalli et al. (2021), we normalized the value of each parameter to facilitate model convergence. We also calculated each participant’s accuracy in two conditions (positive and negative emotional words) to assess whether they have learned the correct rules.

Because previous research has shown that models with lower BIC values indicate a better fit (Liu et al., 2023; Symonds & Moussalli, 2011; Vrieze, 2012), we selected the optimal model according to this criterion. After doing so, if the interaction between variable and parameter was significant, the specific effect of the parameter on the two conditions (positive and negative emotional words) was further examined.

### 2.7 EEG analysis

#### 2.7.1 EEG Recording and Preprocessing

EEG was recorded using a 64-channel Brain products setup, and all electrodes were arranged in accordance with the extended international 10-20 system. The impedance for all electrodes was checked and reduced until it was below 5 kΩ. The FCz electrode was used as the reference electrode while recording, and the average value of TP9 and TP10 amplitudes of the left and right mastoid process electrodes was used for re-reference during offline analysis. EEG activity was filtered with a highpass filter of .1 Hz and a lowpass filter of 30 Hz, with a sampling rate of 500 Hz. Finally, 45 electrodes were left after removing the peripheral electrodes with more artifacts (FP1, FP2, F7, F8, T7, T8, FT9, FT10, AF3, AF4, AF7, AF8, FT7, FT8, TP7, TP8, FPz). All EEG data were analyzed using EEGLAB toolbox (Delorme & Makeig, 2004).

We then used independent component analysis to reduce ocular artifacts. Following this, continuous recordings were epoched around word-locked from −200 to 800 ms. Signals exceeding ± 80 μv were identified and discarded. The number of trials rejected by each subject in each condition did not exceed 8% of the total number of epochs (positive emotional words: 6.82%; negative emotional words: 7.32%).

#### 2.7.2 General linear model regression analysis

To test the modulatory relationship between model parameter and amplitude values, we performed a general linear model regression analysis. Consistent with Hein et al. (2021), for the first-level (subject-specific) analysis, we performed regression analyses for the parameter *μ_2_*. Then, we used the estimated regression coefficients (betas) of the regressor *μ_2_* in each condition to further conduct group-level analyses.

For the second-level (group-specific) inference, the data was performed using the ept-TFCE (threshold-free cluster-enhancement) toolbox (Mensen & Khatami, 2013). This method combined the threshold-free cluster-enhancement technique with permutation-based statistics, which is well suited for mass univariate analyses of event-related potential (ERP) differences between different conditions (Mensen & Khatami, 2013). Given our interest in analyzing the differences of model parameter (*μ_2_*) between the two conditions (positive and negative emotional words), we performed paired *t*-tests at each electrode and each time point (alpha level = .05, number of permutations = 5000) (Kopp et al., 2020).

## 3. Results

### 3.1 Model comparison and selection

We fitted each model (3-level Hierarchical Gaussian Filter [HGF], the Rescorla Wagner [RW], and Sutton K1 [SK1]) with the response data of the 22 individuals and used Bayesian model selection (BMS) to determine the optimal model. Results revealed that the HGF model outperformed the other two models in both conditions because it had the highest expected posterior probabilities, exceedance probabilities, and protected exceedance probabilities (see Table 2). Thus, we chose the HGF model as the best model for the subsequent analyses. Figure 3 shows the three levels of the HGF for binary outcomes and the associated belief trajectories across all 100 trials of positive words for a representative participant.

**Figure 3.**
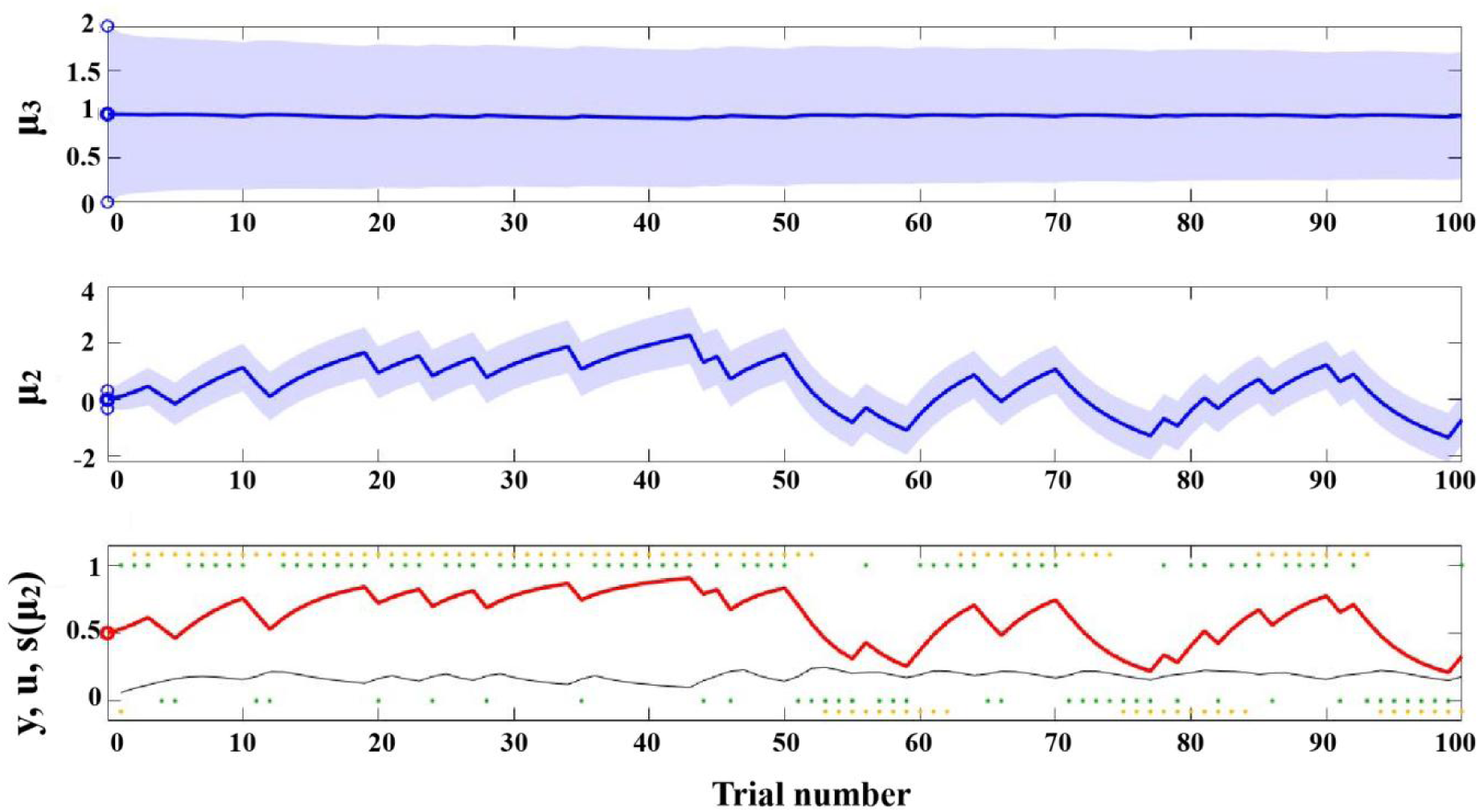
The results of a representative participant. *Notes.* At the lowest level, the inputs *u* (green dots) corresponds to the type of word that was rewarded in each trial (e.g., 1 = words formed with the cu/da suffix, 0 = words formed with the zo/ye suffix), and *y* (orange dots) displays the participant’s responses. The black line and red line represent the participant’s learning rate and posterior expectation of input s(*μ_2_*), respectively. At the second level, the blue line of *μ_2_* represents the participant’s estimate of the picture-word tendency х_2_. At the third level, *μ_2_* (blue line) shows the participant’s estimate of volatility х_3_.

**Table 2.**
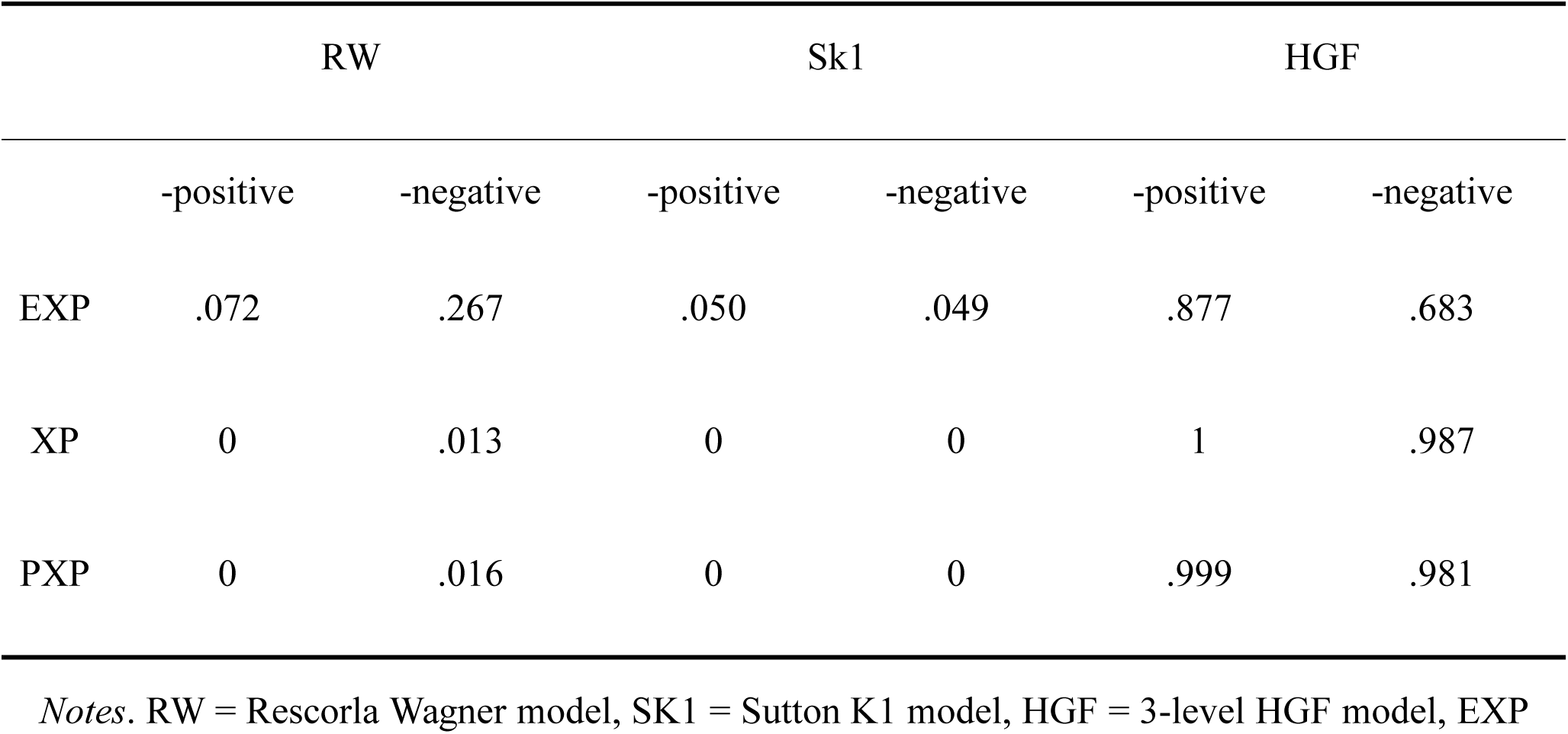
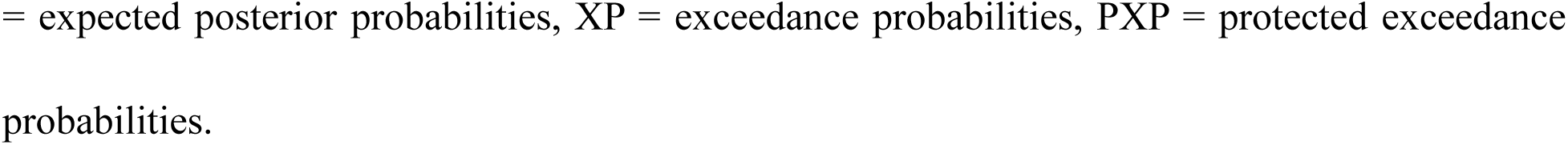
Results of the Bayesian model comparison and selection.

|  | RW |  | Sk1 |  | HGF |  |
| --- | --- | --- | --- | --- | --- | --- |
|  | -positive | -negative | -positive | -negative | -positive | -negative |
| EXP | .072 | .267 | .050 | .049 | .877 | .683 |
| XP | 0 | .013 | 0 | 0 | 1 | .987 |
| PXP | 0 | .016 | 0 | 0 | .999 | .981 |

*Notes.* RW = Rescorla Wagner model, SK1 = Sutton K1 model, HGF = 3-level HGF model, EXP
= expected posterior probabilities, XP = exceedance probabilities, PXP = protected exceedance probabilities.

### 3.2 The impact of HGF on the accuracy and speed of learning words

#### 3.2.1 Accuracy

In the two blocks, the averaged accuracy was 64.591% ± .118% (mean ± standard deviation) in positive emotional word conditions and 69.091% ± .099% in negative emotional word conditions, indicating that participants’ performance was higher than randomized level. In addition, response accuracy under negative emotional words was significantly higher than that under positive emotional words (*t* = 3.204, *p* = .004).

We performed a generalized logistic mixed-effects model on accuracy using word type (positive and negative emotional words) and parameter *μ_2_* as fixed effects. Subjects were added as fixed effects. However, we did not observe a significant interaction between the two variables, which may suggest that *μ_2_* does not have a significant modulatory effect on the accuracy.

#### 3.2.2 RTs

We performed a linear mixed-effects model on log-RTs using word type (positive and negative emotional words) and parameter *μ_2_*. Subjects were added as fixed effects. The results fo the optimal model showed a significant interaction between two variables (*Est* = .04, *SE* = .02, *t* = 2.39, *p* = .02). Further analyses revealed that the model with parameter always outperformed the model without parameter in both conditions, which illustrates the importance of parameter in experiment conditions (see Table 3). As shown in Figure 4, the RTs decreased with increasing *z*-score *μ_2_*, with RTs in the negative word conditions decreasing more rapidly (*Est* = .04, *SE* = .02, *t* = 2.39, *p* = .02).

**Figure 4.**
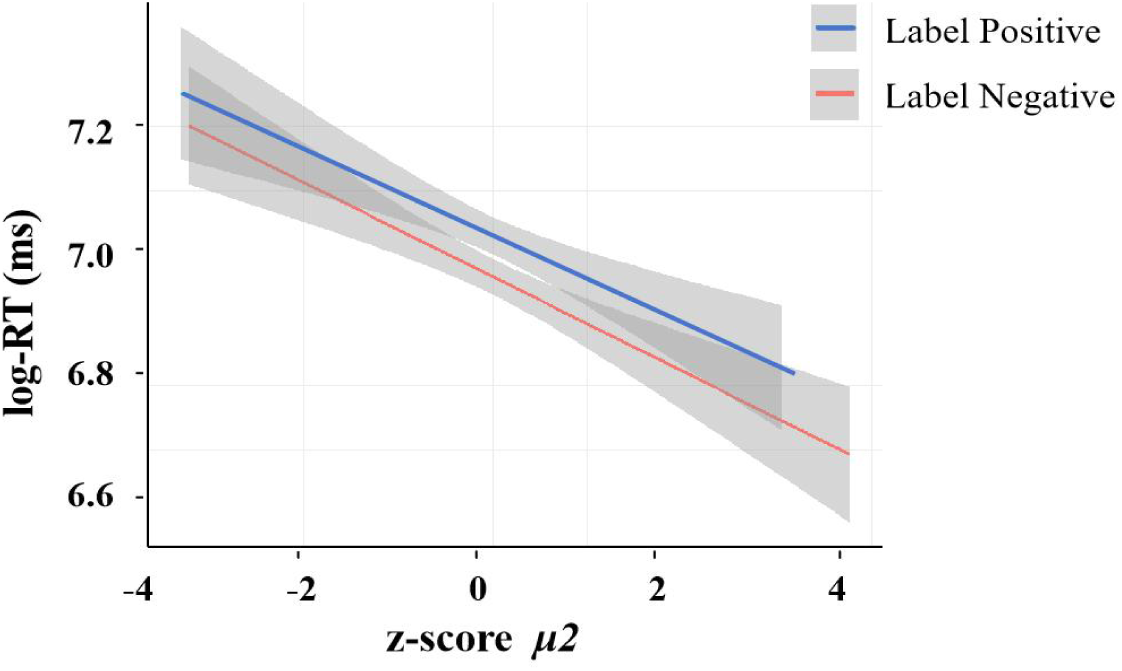
The modulatory effect of *z*-score *μ_2_* on the speed of learning emotional words. *Note.* Grey shading indicates the range of standard deviations for each condition.

**Table 3.** Comparison between models with and without parameter under the positive and negative emotional word conditions.

|  |  | Fixed effect | Random effect | AIC | BIC | Pr ( > chisq) |
| --- | --- | --- | --- | --- | --- | --- |
| Positive | Model-full | $z\_ \mu_2$ | Subject | 3317.9 | 3340.7 | .00073 |
|  | Model-reduced |  | Subject | 3327.3 | 3344.4 |  |
| Negative | Model-full | $z\_ \mu_2$ | Subject | 2872.4 | 2895.2 | .03784 |
|  | Model-reduced |  | Subject | 2874.7 | 2891.8 |  |

### 3.3 Electrophysiological results

#### 3.3.1 Differences between positive and negative emotional words without modulation

The results showed no significant difference between the two conditions (positive and negative emotional words) on the raster diagram, trave-plot, or topoplot (see Figure 5). Therefore, we decided to further investigate whether there were differences between the two conditions under the modulation of *μ_2_*.

**Figure 5.**
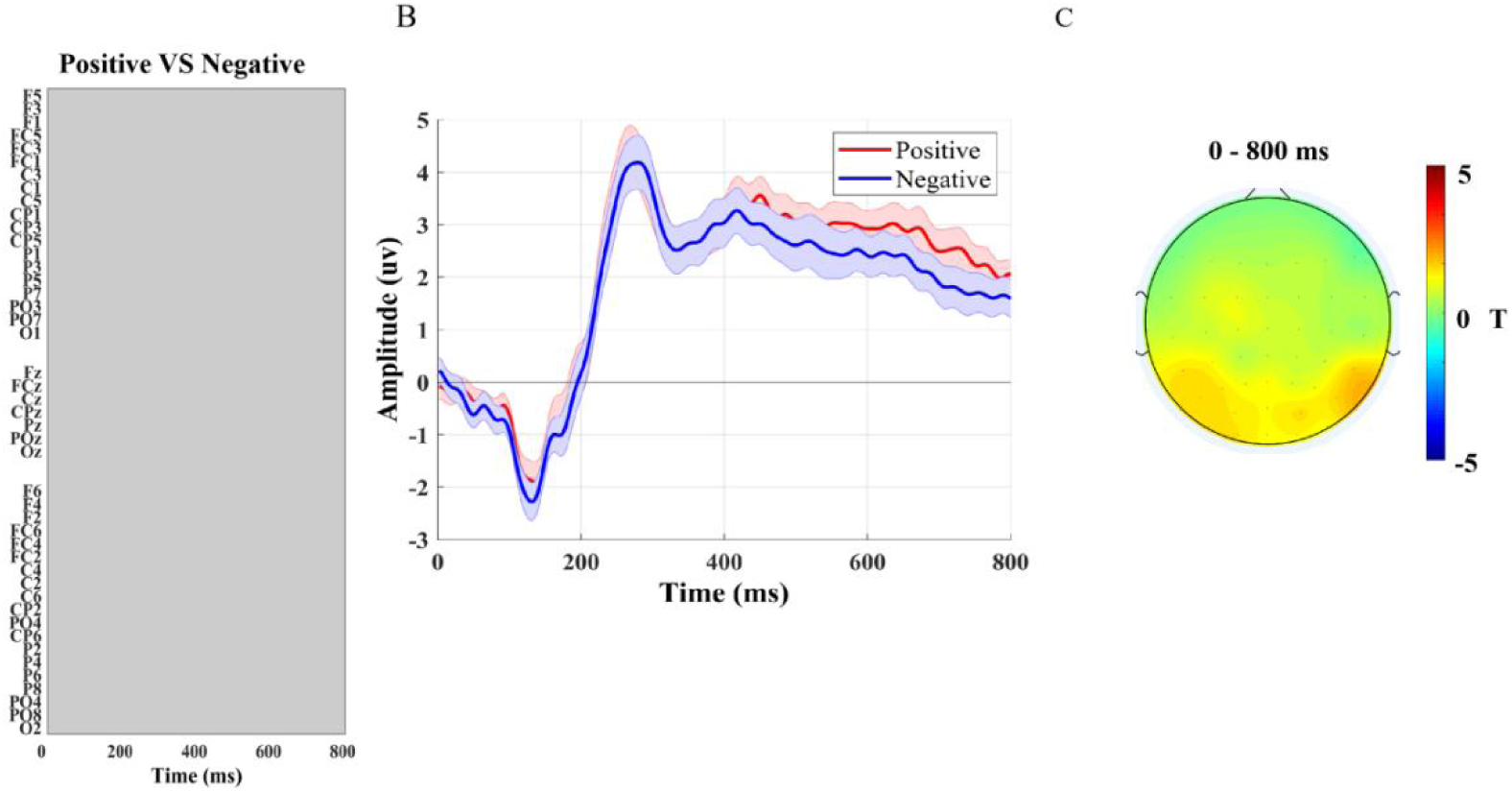
Difference between positive and negative emotional words under unparameterized modulation. A) In the raster diagram, colored rectangles indicate significant differences for different conditions, and grey rectangles indicate no significant difference on any electrodes/time points. In agreement with previous studies (Groppe et al., 2011; Visalli et al., 2021), we display electrodes along the y-axis. B) The trave-plot shows the amplitude of the two conditions (positive and negative emotional words), which are surrounded by standard error bounds. C) The topoplot displays the *t* values averaged for the time window in the diagram (0-800ms).

#### 3.3.2 The modulations of *μ_2_* between positive and negative emotional words

Figure 4 shows the results of the TFCE analysis on the modulatory effect between the two conditions (positive and negative emotional words).

Concerning *μ_2_*, a lasting significant effect was revealed, in which the effect on negative words was higher than the effect on positive words over the right frontal-central electrodes between 270-716 ms. These results are illustrated in Figure 6.

**Figure 6.**
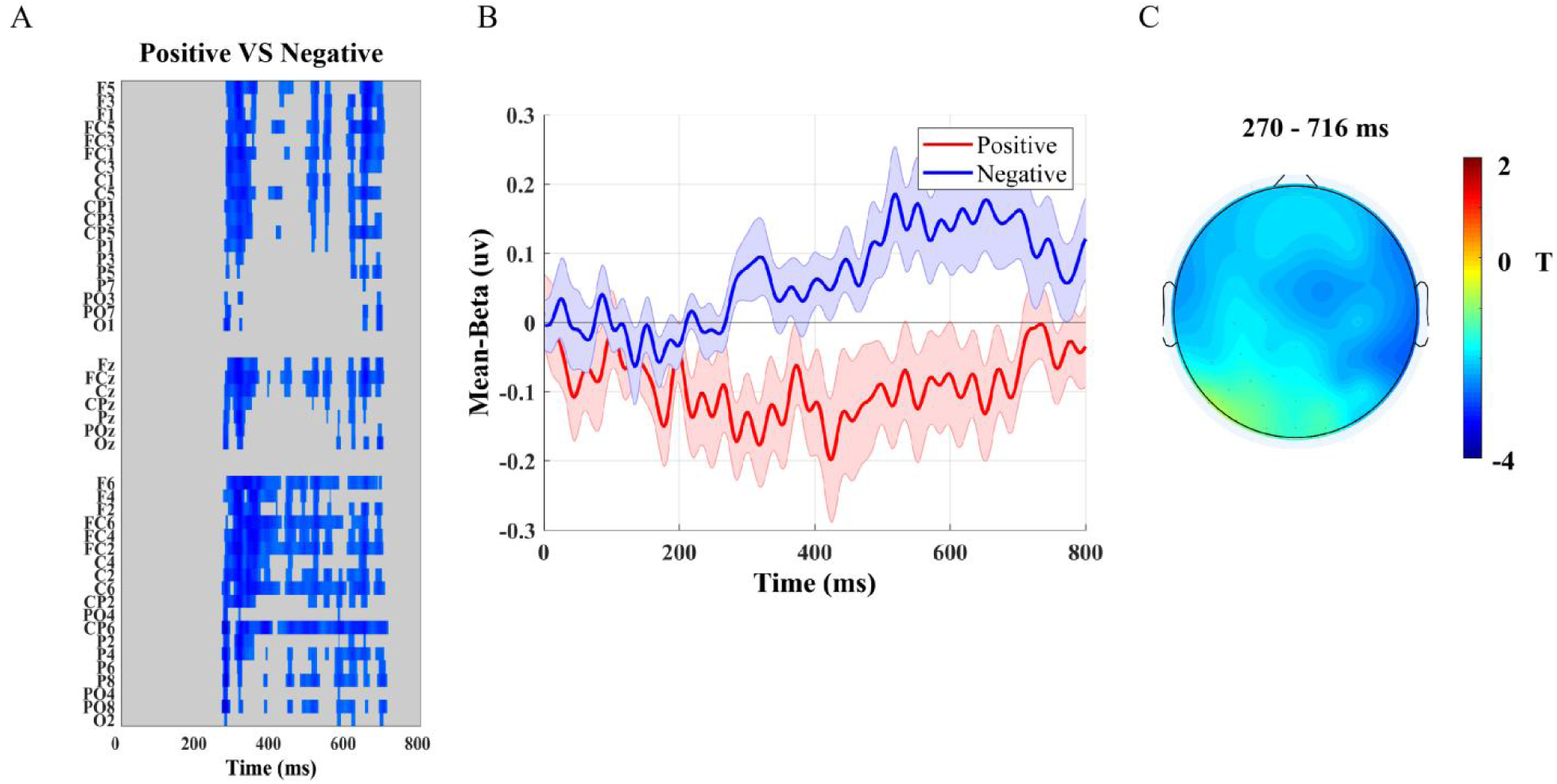
Modulatory differences of the parameter (*μ_2_*) between emotional word type (positive and negative emotional words). A) The colored rectangles in the raster diagram indicate significant differences in the moderating effects of parameters for the different conditions, and grey rectangles indicate no significant moderating effect on any electrodes/time points. B) The trave-plot shows the mean beta values of *μ_2_*, which are surrounded by standard error bounds. C) The topoplot displays the *t* values averaged for the time window in the diagram (270-716ms), which corresponds to the time window of the significant difference in the modulatory effect.

## 4. Discussion

The current study explored how individuals acquire the valence of emotional words in a changing environment by analyzing behavioral and electrophysiological data. The results indicated that the HGF model provided a better fit for the behavioral data. Participants exhibited higher accuracy for negative emotional words, and as *μ_2_* increases, the rate of decrease in RTs under this condition was faster. The EEG results showed that *μ_2_* has a greater modulatory effect on negative emotional words, and the differences between the two conditions (positive and negative emotional words) were concentrated in the right frontal-central region from 270-716 ms. These findings indicate that expectations at the second level (*μ_2_*) modulate the response speed and time course of emotional words learning.

### 4.1 The impact of *μ_2_* on the speed of emotional word learning

In this study, compared to the RW and Sutton models, the HGF demonstrates a clear advantage in fitting behavioral data, which is consistent with previous results (Diaconescu et al., 2014; Hein et al., 2021; Hein & Ruiz, 2022; Iglesias et al., 2013; Iglesias et al., 2021; Liu et al., 2022; Mathys et al., 2014). This finding suggests that individuals may be more inclined to engage in hierarchical learning in a dynamic environment. Our results provide additional behavioral evidence supporting the negative bias in the acquisition of emotional words, consistent with prior studies (Estes & Verges, 2008; Nasrallah et al., 2009; Wu et al., 2022; Zhang et al., 2017). In terms of accuracy, we found that participants were able to complete the experimental task well, and their accuracy was higher under negative emotional words. This indicates that participants were able to discern word formation rules, flexibly perceive changes in the environment, and adjust their choices. Specifically, participants performed better under negative emotional words, which preliminarily indicates a preference for processing negative emotional words.

More importantly, we found that *μ_2_* can modulate an individual’s RTs under two different conditions, and can also exhibit a negativity bias. According to the HGF model, *μ_2_* represents an individual’s expectation about the probability of receiving a reward for given choices (Mathys et al., 2011; Mathys et al., 2014). In the current research, *μ_2_* represented an individual’s expectation of receiving reward for correct emotional words. The results revealed an advantage in processing negative emotional words. Compared to positive emotional words, as *μ_2_* increases, participants responded more quickly to negative emotional words. In sum, the results, whether in terms of RT or accuracy, are consistent with previous research, demonstrating a preference for processing negative emotional words (Estes & Verges, 2008; Nasrallah et al., 2009; Wu et al., 2022; Zhang et al., 2017).

*μ_2_* modulates the temporal dynamics of processing negative emotional words Previous research has shown that processing learning signals can be reflected in ERP amplitudes (Hein et al., 2021; Jepma et al., 2016; Nassar et al., 2019; Liu et al., 2022). Our study has shown that the effect of *μ_2_* on negative emotional words was mainly higher than the effect on positive emotional words over the right frontal-central electrodes between 270-716 ms. This finding is consistent with the behavioral results, both of which demonstrate that *μ_2_* exerts a greater modulatory effect on negative emotional words. On the one hand, this result indicates that the expected value of receiving rewards for certain types of words indeed influences individuals’ processing of those words, thus reflecting the important role of individuals’ expectations in cognition and behavior. In our study, the expectations not only modulated individuals’ RTs, but also the time course of processing. This is consistent with previous research findings that expectations play a crucial role in individuals’ learning and adaptation (Hein et al., 2023; Hein & Ruiz, 2022). On the other hand, during the process of acquiring emotional words, individuals exhibited a clear negative bias: Compared to positive emotional words, when processing negative emotional words, as expectations increased, individuals reacted more rapidly and demonstrated a higher level of neural activity. This provides evidence for the positivity-negativity asymmetry hypothesis (Robinson-Riegler & Winton, 1996), suggesting that due to the greater emotional intensity of negative words, individuals may be more inclined to process the information. The finding also enriches the automatic vigilance hypothesis (Pratto & John, 1991), as negative emotional words may contain threatening information that is closely related to individuals’ survival, and therefore, they may be more inclined to prioritize attention and processing of such information to avoid adverse outcomes.

### 4.2 Limitations and future work

There are some limitations in our study that should be mentioned. First, we used pseudowords as experimental materials and found that individuals’ expectations at the second level (*μ_2_*) significantly modulated their learning of emotional words, particularly demonstrating a preference towards processing negative emotional words. In the future, similar research using real words could be conducted to validate the findings of this study. Second, although our study created a dynamic and changing environment through different probability settings, the degree of fluctuation may have been relatively low. In subsequent work, a more volatile environment should be constructed to explore whether the modulatory effect of expectation values remains stable and whether individuals still exhibit a preference for processing negative emotional words.

Last, previous studies have shown that emotional words encompass both label words and laden words, and that there are differences between them in terms of diverse aspects (Kazanas & Altarriba, 2015, 2016; Knickerbocker & Altarriba, 2013; Knickerbocker et al., 2015; Pavlenko, 2008; Wu & Zhang, 2019; Zhang et al., 2019, 2020). However, the current study only included label words in the experimental materials. In the future, laden words should also be included to provide a more comprehensive explanation for the modulatory effect of expectations on the processing of emotional words.

## 5. Conclusion

This study combines reinforcement learning with emotional word acquisition to reveal how individuals learn the valence of emotional words in stable and volatile environments. The findings of the behavioral and ERP analyses both demonstrated that *μ_2_* can moderate individuals’ preference for processing negative words. Specifically, as *μ_2_* increases, individuals react faster and have greater ERP amplitudes. In summary, our study demonstrates that an individual’s expectations can influence the learning of emotional words in a dynamically changing environment.

## CRediT authorship contribution statement

Weiwei Zhang: Writing – original draft, Methodology, Investigation, Formal analysis, Visualization. Yingyu Li: Methodology, Formal analysis. Chuan Zhou: Investigation, Methodology. Baike Li: Methodology, Formal analysis. John W. Schwieter: Writing–review & editing. Huanhuan Liu: Conceptualization, Methodology, Writing – review & editing, Supervision, Visualization. Meng Liu: Conceptualization, Methodology, Supervision.

## Declaration of Competing Interest

No potential conflict of interest was reported by the authors.

## Data availability

Data will be made available on request.

## Supplementary Materials

**Table S1.** Pseudowords under positive and negative conditions.

| Positive emotional words |  |  |  |  |  |
| --- | --- | --- | --- | --- | --- |
| Stable | necu | sacu | pucu | kicu | hocu |
|  | tuzo | gizo | qezo | mozo | razo |
| Volatile | nezo | sazo | puzo | kizo | hozo |
|  | tucu | gicu | qecu | mocu | racu |
| Negative emotional words |  |  |  |  |  |
| Stable | neda | sada | puda | kida | hoda |
|  | tuye | giye | qeye | moye | raye |
| Volatile | neye | saye | puye | kiye | hoye |
|  | tuda | gida | qeda | moda | rada |

## Notes

**Author Notes** This work was supported by a Grant from STI 2030—Major Projects 2021ZD0200500, General Program of National Natural Science Foundation of China (32371089), Liaoning Social Science Planning Fund of China (L20AYY001), Dalian Science and Technology Star Fund of China (2020RQ055), and the Youth Project of Liaoning Provincial Department of Education (LJKQZ2021089), Research and Cooperation Projects on Social and Economic Development of Liaoning Province (2024lslybhzkt-17), and Liaoning Educational Science Planning Project (JG21DB306). We have no known competing interests to declare.

### Competing Interest Statement

The authors have declared no competing interest.

## References

Albrecht, C., van de Vijver, R., & Bellebaum, C. (2023). Learning new words via feedback—Association between feedback-locked ERPs and recall performance—An exploratory study. Psychophysiology, 60(10), e14324.

Altarriba, J., & Basnight-Brown, D. M. (2011). The representation of emotion vs. emotion-laden words in English and Spanish in the Affective Simon Task. International Journal of Bilingualism, 15(3), 310–328.

Bayer, M., & Schacht, A. (2014). Event-related brain responses to emotional words, pictures, and faces–a cross-domain comparison. Frontiers in Psychology, 5, 1106.

Browning, M., Behrens, T. E., Jocham, G., O’reilly, J. X., & Bishop, S. J. (2015). Anxious individuals have difficulty learning the causal statistics of aversive environments. Nature Neuroscience, 18(4), 590–596.

Damasio, A. (2001). Fundamental feelings. Nature, 413(6858), 781–781.

De Berker, A. O., Rutledge, R. B., Mathys, C., Marshall, L., Cross, G. F., Dolan, R. J., & Bestmann, S. (2016). Computations of uncertainty mediate acute stress responses in humans. Nature Communications, 7(1), 10996.

Delorme, A., & Makeig, S. (2004). EEGLAB: An open-source toolbox for analysis of single-trial EEG dynamics including independent component analysis. Journal of Neuroscience Methods, 134(1), 9–21.

Degner, J., Doycheva, C., & Wentura, D. (2012). It matters how much you talk: On the automaticity of affective connotations of first and second language words. Bilingualism: Language and Cognition, 15(1), 181–189.

Den Ouden, H. E., Daw, N. D., Fernandez, G., Elshout, J. A., Rijpkema, M., Hoogman, M.,… & Cools, R. (2013). Dissociable effects of dopamine and serotonin on reversal learning. Neuron, 80(4), 1090–1100.

Diaconescu, A. O., Mathys, C., Weber, L. A., Daunizeau, J., Kasper, L., Lomakina, E. I.,… & Stephan, K. E. (2014). Inferring on the intentions of others by hierarchical Bayesian learning. PLoS Computational Biology, 10(9), e1003810.

Douglas Bates, M. M., Bolker, B., & Walker, S. (2015). Fitting linear mixed-effects models using lme4. Journal of Statistical Software, 67(1), 1–48.

Eilola, T. M., Havelka, J., & Sharma, D. (2007). Emotional activation in the first and second language. Cognition and Emotion, 21(5), 1064–1076.

Eilola, T. M., & Havelka, J. (2011). Behavioural and physiological responses to the emotional and taboo Stroop tasks in native and non-native speakers of English. International Journal of Bilingualism, 15(3), 353–369.

Estes, Z., & Verges, M. (2008). Freeze or flee? Negative stimuli elicit selective responding. Cognition, 108(2), 557–565.

Frishkoff, G. A., Collins-Thompson, K., Hodges, L., & Crossley, S. (2016). Accuracy feedback improves word learning from context: Evidence from a meaning-generation task. Reading and Writing, 29, 609–632.

Groppe, D. M., Urbach, T. P., & Kutas, M. (2011). Mass univariate analysis of event related brain potentials/fields I: A critical tutorial review. Psychophysiology,48 (12), 1711–1725.

Goldstein, M. H., & Schwade, J. A. (2008). Social feedback to infants’ babbling facilitates rapid phonological learning. Psychological Science, 19(5), 515–523.

Gong, T., Lam, Y. W., & Shuai, L. (2016). Influence of perceptual saliency hierarchy on learning of language structures: an artificial language learning experiment. Frontiers in Psychology, 7, 233336.

Hackel, L., & Amodio, D. (2018). Computational neuroscience approaches to social cognition. Current Opinion in Psychology, 2018, 24, 92–97.

Hackel, L. M., Mende-Siedlecki, P., & Amodio, D. M. (2020). Reinforcement learning in social interaction: The distinguishing role of trait inference. Journal of Experimental Social Psychology, 88, 103948.

Held, L. K., Vermeylen, L., Dignath, D., Notebaert, W., Krebs, R. M., & Braem, S. (2024). Reinforcement learning of adaptive control strategies. Communications Psychology, 2(1), 8.

Hein, T. P., de Fockert, J., & Ruiz, M. H. (2021). State anxiety biases estimates of uncertainty and impairs reward learning in volatile environments. NeuroImage, 224, 117424.

Hein, T. P., & Ruiz, M. H. (2022). State anxiety alters the neural oscillatory correlates of predictions and prediction errors during reward-based learning. NeuroImage, 249, 118895.

Hein, T. P., Gong, Z., Ivanova, M., Fedele, T., Nikulin, V., & Herrojo Ruiz, M. (2023). Anterior cingulate and medial prefrontal cortex oscillations underlie learning alterations in trait anxiety in humans. Communications Biology, 6(1), 271.

Hinojosa, J. A., Mercado, F., Albert, J., Barjola, P., Peláez, I., Villalba-García1, C., & Carretié, L. (2015). Neural correlates of an early attentional capture by positive distractor words. Frontiers in Psychology, 6(1), 1–13.

Iglesias, S., Kasper, L., Harrison, S. J., Manka, R., Mathys, C., & Stephan, K. E. (2021). Cholinergic and dopaminergic effects on prediction error and uncertainty responses during sensory associative learning. NeuroImage, 226, 117590.

Iglesias, S., Mathys, C., Brodersen, K. H., Kasper, L., Piccirelli, M., den Ouden, H. E., & Stephan, K. E. (2013). Hierarchical prediction errors in midbrain and basal forebrain during sensory learning. Neuron, 80(2), 519–530.

Jepma, M., Murphy, P. R., Nassar, M. R., Rangel-Gomez, M., Meeter, M., & Nieuwenhuis, S. (2016). Catecholaminergic regulation of learning rate in a dynamic environment. PLoS Computational Biology, 12(10), e1005171.

Kauschke, C., Bahn, D., Vesker, M., & Schwarzer, G. (2019). The role of emotional valence for the processing of facial and verbal stimuli: Positivity or negativity bias? Frontiers in Psychology, 10, 1654.

Kazanas, S. A., & Altarriba, J. (2015). The automatic activation of emotion and emotion-laden words: Evidence from a masked and unmasked priming paradigm. The American Journal of Psychology, 128(3), 323–336.

Kazanas, S. A., & Altarriba, J. (2016). Emotion word processing: Effects of word type and valence in Spanish-English bilinguals. Journal of Psycholinguistic Research, 45(2), 395–406

Knickerbocker, H., & Altarriba, J. (2013). Differential repetition blindness with emotion and emotion-laden word types. Visual Cognition, 21(5), 599–627.

Knickerbocker, H., Johnson, R. L., & Altarriba, J. (2015). Emotion effects during reading: Influence of an emotion target word on eye movements and processing. Cognition and Emotion, 29(5), 784–806.

Kopp, B., Steinke, A., & Visalli, A. (2020). Cognitive flexibility and N2/P3 event-related brain potentials. Scientific Reports, 10(1), 9859.

Kuperman, V., Estes, Z., Brysbaert, M., & Warriner, A. B. (2014). Emotion and language: valence and arousal affect word recognition. Journal of Experimental Psychology: General, 143(3), 1065.

Lee, D., Seo, H., & Jung, M. W. (2012). Neural basis of reinforcement learning and decision making. Annual Review of Neuroscience, 35(1), 287–308.

Lefebvre, G., Lebreton, M., Meyniel, F., Bourgeois-Gironde, S., & Palminteri, S. (2017). Behavioural and neural characterization of optimistic reinforcement learning. Nature Human Behaviour, 1(4), 0067.

Liu, M., Dong, W., Qin, S., Verguts, T., & Chen, Q. (2022). Electrophysiological signatures of hierarchical learning. Cerebral Cortex, 32(3), 626–639.

Liu, L., Liu, D., Guo, T., Schwieter, J. W., & Liu, H. (2023). The right superior temporal gyrus plays a role in semantic-rule learning: Evidence supporting a reinforcement learning model. NeuroImage, 282, 120393.

Li, S., Seger, C. A., Zhang, J., Liu, M., Dong, W., Liu, W., & Chen, Q. (2023). Alpha oscillations encode Bayesian belief updating underlying attentional allocation in dynamic environments. NeuroImage, 284, 120464.

Lockwood, P. L., & Klein-Flügge, M. C. (2021). Computational modelling of social cognition and behaviour: A reinforcement learning primer. Social Cognitive and Affective Neuroscience, 16(8), 761–771.

Mathys, C., Daunizeau, J., Friston, K. J., & Stephan, K. E. (2011). A Bayesian foundation for individual learning under uncertainty. Frontiers in Human Neuroscience, 5, 39.

Mathys, C. D., Lomakina, E. I., Daunizeau, J., Iglesias, S., Brodersen, K. H., Friston, K. J., & Stephan, K. E. (2014). Uncertainty in perception and the Hierarchical Gaussian Filter. Frontiers in Human Neuroscience, 8, 825.

Mensen, A., & Khatami, R. (2013). Advanced EEG analysis using threshold-free cluster-enhancement and non-parametric statistics. NeuroImage, 67, 111–118.

Metha, J. A., Brian, M. L., Oberrauch, S., Barnes, S. A., Featherby, T. J., Bossaerts, P.,… & Jacobson, L. H. (2020). Separating probability and reversal learning in a novel probabilistic reversal learning task for mice. Frontiers in Behavioral Neuroscience, 13, 270.

Mukherjee, D., Filipowicz, A. L., Vo, K., Satterthwaite, T. D., & Kable, J. W. (2020). Reward and punishment reversal-learning in major depressive disorder. Journal of Abnormal Psychology, 129(8), 810.

Nasrallah, M., Carmel, D., & Lavie, N. (2009). Murder, she wrote: Enhanced sensitivity to negative word valence. Emotion, 9(5), 609.

Nassar, M. R., McGuire, J. T., Ritz, H., & Kable, J. W. (2019). Dissociable forms of uncertainty-driven representational change across the human brain. Journal of Neuroscience, 39(9), 1688–1698.

Palminteri, S., & Lebreton, M. (2022). The computational roots of positivity and confirmation biases in reinforcement learning. Trends in Cognitive Sciences, 26(7), 607–621.

Pashler, H., Cepeda, N. J., Wixted, J. T., & Rohrer, D. (2005). When does feedback facilitate learning of words? Journal of Experimental Psychology: Learning, Memory, and Cognition, 31(1), 3.

Pavlenko, A. (2008). Emotion and emotion-laden words in the bilingual lexicon. Bilingualism: Language and Cognition, 11(2), 147–164.

Ponari, M., Rodríguez-Cuadrado, S., Vinson, D., Fox, N., Costa, A., & Vigliocco, G. (2015). Processing advantage for emotional words in bilingual speakers. Emotion, 15(5), 644.

Pons, F., & Harris, P. (2000). Test of emotion comprehension–TEC. Oxford University Press.

Pons, F., Harris, P. L., & de Rosnay, M. (2004). Emotion comprehension between 3 and 11 years: Developmental periods and hierarchical organizations. European Journal of Developmental Psychology, 1, 127–152.

Pratto, F., & John, O. P. (1991). Automatic vigilance: The attention-grabbing power of negative social information. Journal of Personality and Social Psychology, 61(3), 380.

Rescorla, R. A., & Wagner, A. R. (1972). A theory of Pavlovian conditioning: Variations in the effectiveness of reinforcement and nonreinforcement. Appleton-Century-Crofts.

Robinson-Riegler, G. L., & Winton, W. M. (1996). The role of conscious recollection in recognition of affective material: Evidence for positive-negative asymmetry. The Journal of General Psychology, 123(2), 93–104.

Russell, J. A. (1980). A circumplex model of affect. Journal of Personality and Social Psychology, 39(6), 1161–1178.

Schindler, S., & Kissler, J. (2016). Selective visual attention to emotional words: Early parallel frontal and visual activations followed by interactive effects in visual cortex. Human Brain Mapping, 37(10), 3575–3587.

Schindler, S., Vormbrock, R., & Kissler, J. (2019). Emotion in context: how sender predictability and identity affect processing of words as imminent personality feedback. Frontiers in Psychology, 10, 94.

Scott, G. G., O’Donnell, P. J., Leuthold, H., & Sereno, S. C. (2009). Early emotion word processing: Evidence from event-related potentials. Biological Psychology, 80(1), 95–104.

Sharp, P. B., Russek, E. M., Huys, Q. J., Dolan, R. J., & Eldar, E. (2022). Humans perseverate on punishment avoidance goals in multigoal reinforcement learning. Elife, 11, e74402.

Smith, N. K., Cacioppo, J. T., Larsen, J. T., & Chartrand, T. L. (2003). May I have your attention, please: Electrocortical responses to positive and negative stimuli. Neuropsychologia, 41(2), 171–183.

Snefjella, B., Lana, N., & Kuperman, V. (2020). How emotion is learned: Semantic learning of novel words in emotional contexts. Journal of Memory and Language, 115, 104171.

Soch, J., Haynes, J. D., & Allefeld, C. (2016). How to avoid mismodelling in GLM-based fMRI data analysis: Cross-validated Bayesian model selection. NeuroImage, 141, 469–489.

Stephan, K. E., Penny, W. D., Daunizeau, J., Moran, R. J., & Friston, K. J. (2009). Bayesian model selection for group studies. NeuroImage, 46(4), 1004–1017.

Sutton, R. S. (1992). Gain adaptation beats least squares. Proceedings of the 7^th^ Yale Workshop on Adaptive and Learning Systems, 161, 166.

Sutton, T. M., Altarriba, J., Gianico, J. L., & Basnight-Brown, D. M. (2007). The automatic access of emotion: Emotional Stroop effects in Spanish–English bilingual speakers. Cognition and Emotion, 21(5), 1077–1090.

Sylvester, T., Braun, M., Schmidtke, D., & Jacobs, A. M. (2016). The Berlin affective word list for children (kidBAWL): Exploring processing of affective lexical semantics in the visual and auditory modalities. Frontiers in Psychology, 7, 969.

Symonds, M. R., & Moussalli, A. (2011). A brief guide to model selection, multimodel inference and model averaging in behavioural ecology using Akaike’s information criterion. Behavioral Ecology and Sociobiology, 65, 13–21.

Taylor, S. E. (1991). Asymmetrical effects of positive and negative events: The mobilization-minimization hypothesis. Psychological Bulletin, 110(1), 67.

Unkelbach, C., Fiedler, K., Bayer, M., Stegmüller, M., & Danner, D. (2008). Why positive information is processed faster: The density hypothesis. Journal of Personality and Social Psychology, 95(1), 36–49.

Vrieze, S. I. (2012). Model selection and psychological theory: A discussion of the differences between the Akaike information criterion (AIC) and the Bayesian information criterion (BIC). Psychological Methods, 17(2), 228.

Visalli, A., Capizzi, M., Ambrosini, E., Kopp, B., & Vallesi, A. (2021). Electroencephalographic correlates of temporal Bayesian belief updating and surprise. NeuroImage, 231, 117867.

Wang, Z., Nan, T., Goerlich, K. S., Li, Y., Aleman, A., Luo, Y., & Xu, P. (2023). Neurocomputational mechanisms underlying fear-biased adaptation learning in changing environments. PLoS Biology, 21(5), e3001724.

Wu, C., & Zhang, J. (2019). Conflict processing is modulated by positive emotion word type in second language: An ERP study. Journal of Psycholinguistic Research, 48(5), 1203–1216.

Wu, C., Zhang, J., & Yuan, Z. (2022). An ERP investigation on the second language and emotion perception: The role of emotion word type. International Journal of Bilingual Education and Bilingualism, 25(2), 539–551.

Yao, Z., Yu, D., Wang, L., Zhu, X., Guo, J., & Wang, Z. (2016). Effects of valence and arousal on emotional word processing are modulated by concreteness: Behavioral and ERP evidence from a lexical decision task. International Journal of Psychophysiology, 110, 231–242.

Zhang, J., Teo, T., & Wu, C. (2019). Emotion words modulate early conflict processing in a flanker task: Differentiating emotion-label words and emotion-laden words in second language. Language and Speech, 62(4), 641–651.

Zhang, J., Wu, C., Meng, Y., & Yuan, Z. (2017). Different neural correlates of emotion-label words and emotion laden words: An ERP study. Frontiers in Human Neuroscience, 11(9), 455–463.

Zhang, J., Wu, C., Yuan, Z., & Meng, Y. (2019). Differentiating emotion-label words and emotion-laden words in emotion conflict: An ERP study. Experimental Brain Research, 237(9), 2423–2430.

Zhang, J., Wu, C., Yuan, Z., & Meng, Y. (2020). Different early and late processing of emotion-label words and emotion-laden words in a second language: An ERP study. Second Language Research, 36(3), 399–412.

Zhang, T., Zhou, S., Bai, X., Zhou, F., Zhai, Y., Long, Y., & Lu, C. (2023). Neurocomputations on dual-brain signals underlie interpersonal prediction during a natural conversation. NeuroImage, 282, 120400.

